# Multiple modulation synthesis with high spatial resolution for noninvasive deep neurostimulation

**DOI:** 10.1101/416057

**Authors:** Qiaoqin Xiao, Xiaozheng Lai, Hao Chen

## Abstract

Noninvasive neurostimulation plays a pivotal role in direct control of neural circuit and modulating neuronal function. However, it is difficult to balance both spatial focality and depth of penetration in stimulating deep neurons. Here, we designed time-division, frequency & polarity modulation synthesis (TMFPMS) for stimulating deep neurons noninvasively with low-frequency envelope. We demonstrated its spatial resolution: mm-level via computational simulation including finite element analysis and Hodgkin-Huxley action potential model. Taken together, the results of this study indicate that TMFPMS neurostimulation with high spatial resolution is steerable and may replace traditional implanted electrode.

## 1 INTRODUCTION

Electrical stimulation has been widely used in therapeutic applications, including treatment of neurological and psychiatric disorders through modulating neuronal functions with high spatial and temporal resolution. Nevertheless, vagus nerve stimulation (Groves & Brown, 2005) and deep brain stimulation (DBS) (Montgomery, 2017), with implanted electrode, are accompanied with invasive operations, which may increase hazardous risk.

Non-invasive stimulations, such as transcranial magnetic stimulation (TMS) (Deng, Lisanby & Peterchev, 2013), transcranial alternating current stimulation (tACs) and transcranial direct current stimulation (tDCs) (Datta et al., 2008), produce magnetic fields or electric fields to modulate excitability of neurons through in vitro way. But above of them have low spatial precision (cm-level), compared to the effects of invasive methods (mm-level). Meanwhile, they only directly affect activity in cortical regions and are difficult to stimulate deeper neurons, such as hippocampus or subthalamic nucleus. Besides, transcranial focused ultrasound stimulation (Rezayat & Toostani, 2016) are more focal than both TMS and TES (transcranial electrical stimulation) as its effects are expressed in mm-level. However, its mechanism of action is not clear and may damage other tissues.

Nerve noninvasive electrical stimulation research trends are both in depth and high spatial resolution. For example, concentric electrode (Mücke et al., 2014) and ring electrode (Datta et al., 2009) can provide better spatial focality but need higher applying voltage than bipolar electrode. Apparently, the higher stimulating current, the deeper neurons can be effectively stimulated effectively, but the diffusion of the electric current induces unintended excitation of other neurons and deteriorates the accuracy of the electrical stimulation. Because evaluating the effective stimulation region depends largely on whether current density exceeds the specific threshold current. All of the above mentioned noninvasive methods, there is a trade-off between depth of penetration and spatial focality (Deng, Lisanby & Peterchev, 2013).

Recently, temporally interfering electric field (Grossman et al., 2017) have been investigated in mouse that interferential current can stimulate deep-lying hippocampus without recruitment of overlying cortex. Because in deeper region, temporally interfering electric field induced low-frequency envelope which had enough relaxation period whenever the superimposed waveforms were below current threshold. It can overcome high-frequency fatigue of neurons. At the same time, kilohertz-frequency alternating current, such as burst-modulated and premodulated interferential alternating current (Ward, 2009), can produce depth-efficient stimulation of nerve and muscle thanks to high penetration of kilohertz-frequency in human tissue. Intersectional pulse stimulation (Vöröslakos et al., 2018) distributed currents and regulated focus area via multiple electrodes by temporal multiplexing. However, the spatial precision of interferential current and intersectional pulse neurostimulation did not have a quantification or needed to be improved, perhaps using multiple sets of electric fields.

The goal of this paper is to introduce a novel noninvasive method that can stimulate desired neurons at depth with high spatial resolution. Utilizing multiple electrodes on scalp, applying time-division, frequency & polarity modulation synthesis (TDFPMS) current, can control excitable region of deep neurons to mm-level region.

We utilized finite element method (FEM) to calculate distribution of stimulation waveform in multi-layered model. Additionally, somas were established in NEURON software to evaluate neural excitability across the whole brain. Integrating FEM and NEURON, we concluded that our noninvasive stimulation method can stimulate deep neurons effectively with high spatial resolution: mm-level. TDFPMS method for neurostimulation with high spatial resolution may replace traditional implanted electrodes, such as traditional deep brain stimulation and vagus nerve stimulation.

## 2 MATERIALS AND METHODS

### Time-division, frequency & polarity modulation synthesis (TDFPMS)

The target region (SUM) in response to stimulation waveform with low-frequency standard envelope and high carrier frequency as shown in Fig. 1. It is superimposed by multiple channels (CH 1 – CH 8) based on time-division, frequency & polarity modulation synthesis (TDFPMS). In specific time slot, some channels output different frequency half sine wave that positive and negative polarity were separate. For example, the first group of electrodes (CH 1 – CH 4), with (frequency, polarity) of (*f*_*c*_ + Δ*f*, positive), (*f*_*c*_ + Δ*f*, negative), (*f*_*c*_, positive) and (*f*_*c*_, negative), make up low-frequency envelope in the first time slot. Meanwhile, positive and negative polarity channels at the same frequency superimpose to form a sine wave. Then sum of two sine waves, with different frequencies (*f*_*c*_ + Δ*f* and *f*_*c*_), generate envelope modulated at the low frequency Δ*f*. Similarly, in the second time slot, CH 5 - CH 8 make up the other low-frequency envelope. Temporal summation of several groups of envelopes, generated by electrodes distributed around scalp, creates standard low-frequency envelope in the target region and elicits most neural excitability in complete cycle.

**Figure 1:**
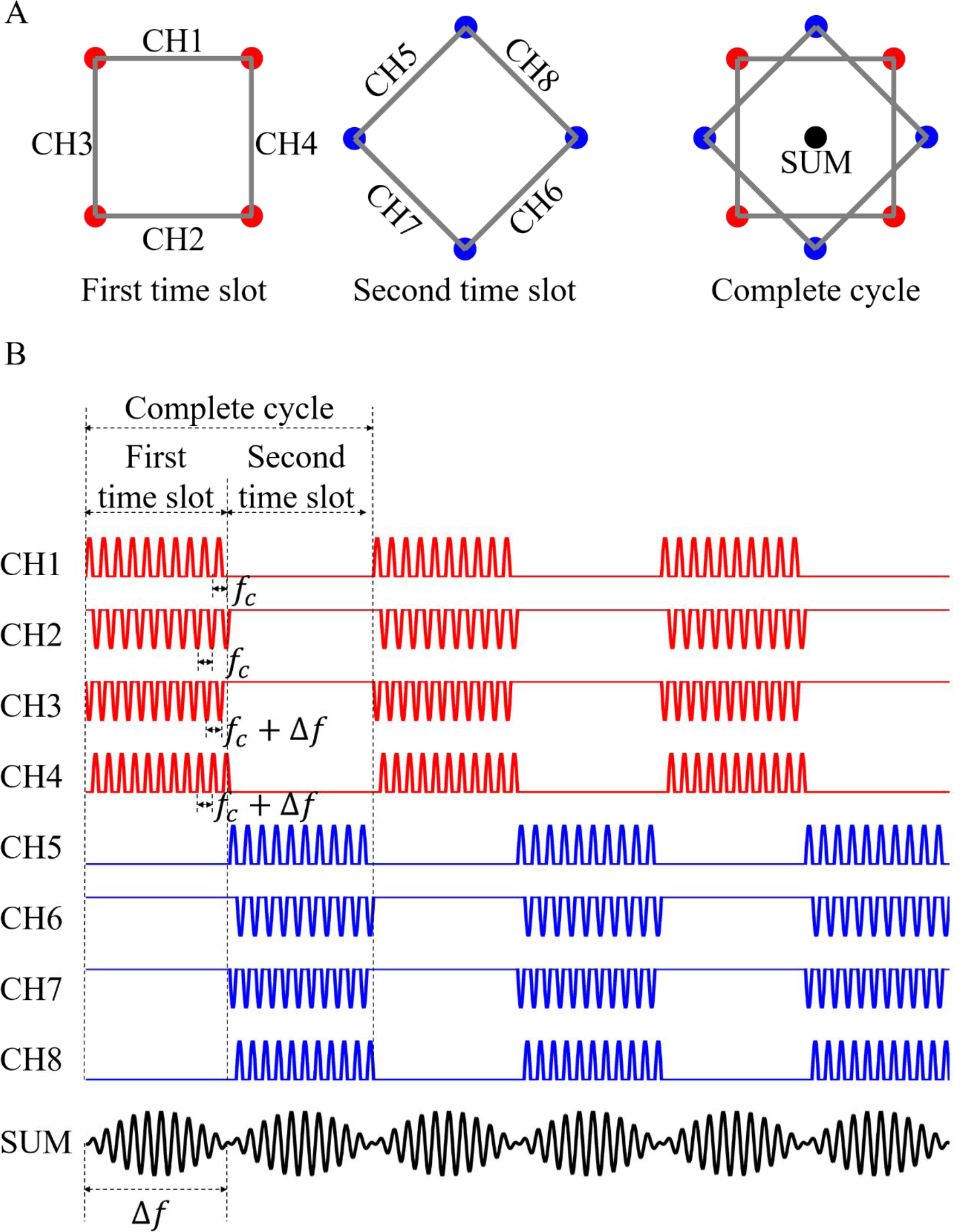
Illustration of Time-Division, Frequency & Polarity Modulation Synthesis (TDFPMS). (A) Arrangements of multiple electrodes acting in the first time slot (red), second time slot (blue) and complete cycle (black) respectively. SUM: target location. (B) Waveforms outputting from various channels and their summation. Please note that the length of kilohertz-half sine wave is shown disproportionally for better visibility.

### FEM with multi-layered tissue and multiple electrodes

Finite element method (FEM) of simplified multi-layered tissue medium and multiple electrode was created to calculate the induced current density generated by each set of electrodes respectively as shown in Fig. 2. The brain model, was simplified into concentric spheres, comprised of four layers: scalp, skull, cerebrospinal fluid (CSF) and brain. Table 1 lists the size, relative permittivity ε_r_ and conductivity σ of above tissue medium (Andreuccetti, Fossi & Petrucci, 2002). Electrodes, with size of 25 mm × 25 mm and thickness of 2mm, were placed around the scalp surface. Each set of electrodes was driven with voltage-source terminal and ground boundary condition. Using frequency domain solution, instead of stationary solution for direct current, calculated various current densities generated by each set of electrodes in the tissue medium respectively.

**Table 1:** Geometrical dimensions and dielectric properties of tissue at 2 kHz frequency

| | Relative permittivity ($\epsilon_r$) | Conductivity $\sigma$ (S/m) | Radical thickness (mm) |
| --- | --- | --- | --- |
| Skin | 31034 | 0.0008 | 2 |
| Skull | 1700 | 0.0202 | 2 |
| CSF | 109 | 2 | 2 |
| Brain | 94300 | 0.1230 | 94 |

**Figure 2:**
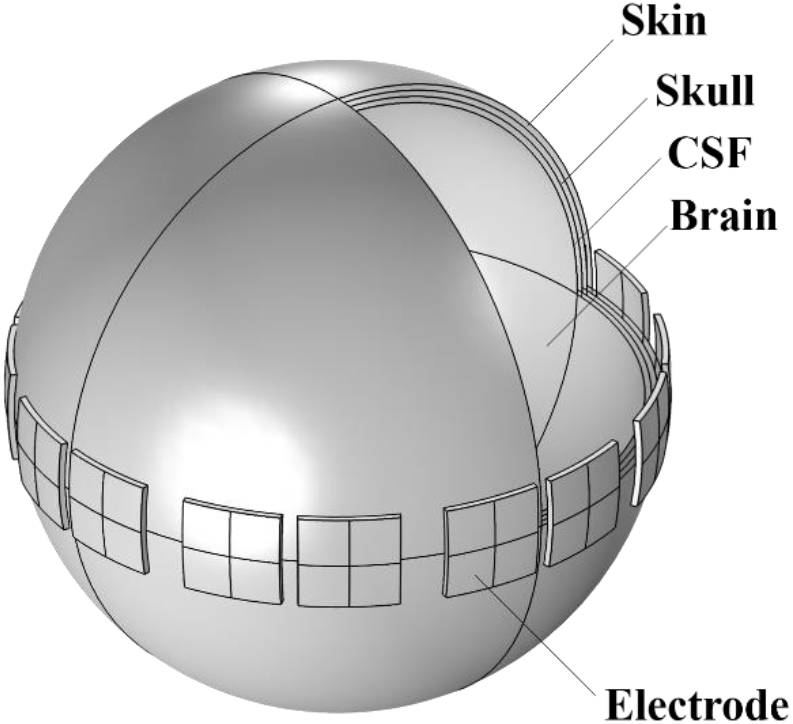
FEM model for noninvasive deep brain stimulation. Include electrode pairs and four tissue layers: scalp, skull, CSF and brain.

The frequency domain solution is based on Maxwell’s equations:

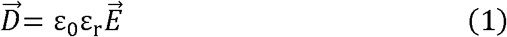

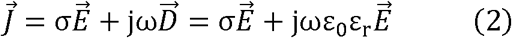

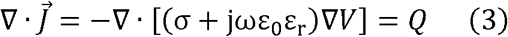

where 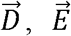 and 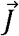 represent displacement field, electric field and electric current density respectively, ω is frequency, ε_r_ is relative permittivity, *Q* is total charge, ∇. is the divergence of a vector function and ∇ is a scalar function. From Equation 2, the former one is conduct current 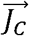 and the latter one is displacement current 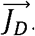. Owning to high relative permittivity of tissue (see Table 1), the displacement current 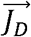 with kilohertz frequency should not be ignored (Stoykov et al., 2002). Therefore, with the same applied voltage, the current density 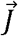 generated by kilohertz-frequency alternating current (AC) in deep tissue is larger than low-frequency AC or direct current. In summary, kilohertz-frequency AC has a good penetration across tissue medium due to its higher displacement, corresponding to lower capacitive resistance 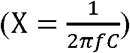 in tissue medium.

### Modeling of Action potentials

We used the well-known Hodgkin-Huxley (H-H) model to establish point neurons and evaluate the effect of specific current waveforms as a current injected into a soma. It takes into account three fundamental active membrane channels: Na^+^ channels, K^+^ channels and leakage channel, consisting of four differential equations:

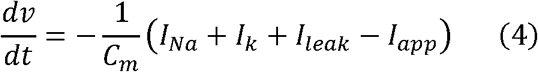

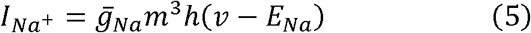

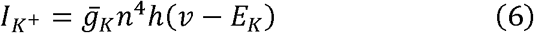

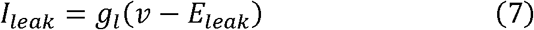

All gating variables are voltage dependent and given by the following equations:

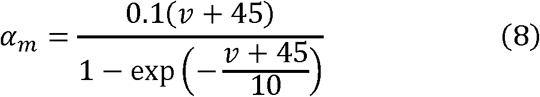

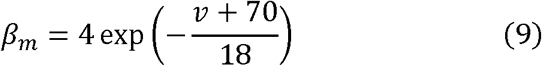

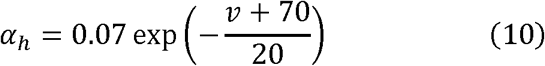

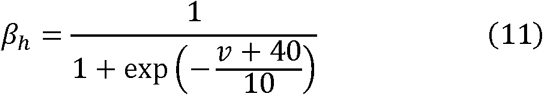

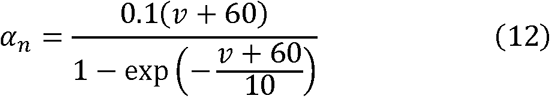

This approach captured the induced membrane potential causing by external injected current at the level of soma. To improve computational efficiency and accuracy, soma (L = 96 µm, D = 96 µm) was constructed in NEURON software (Hines & Carnevale, 1997). All parameters for the soma and Hodgkin–Huxley model are shown in Table 2 (Hoppensteadt & Peskin, 2010). An effective action potential is simply defined as: membrane potential crossed a threshold 30 mV in the absence of fatigue.

**Table 2:** Model parameters for the Hodgkin–Huxley model and soma.

| Name | Value & unit | Description |
| --- | --- | --- |
| $C_m$ | 1 $\mu\text{F}/\text{cm}^2$ | Membrane capacitance |
| $\bar{g}_{Na}$ | 120 $\text{m}/\text{cm}^2$ | Sodium conductance |
| $\bar{g}_K$ | 36 $\text{m}/\text{cm}^2$ | Potassium leak conductance |
| $g_l$ | 0.3 $\text{m}/\text{cm}^2$ | Leak conductance |
| $E_{Na}$ | 45 mV | Sodium reversal potential |
| $E_k$ | -82 mV | Potassium reversal potential |
| $E_{leak}$ | -59 mV | Leak reversal potential |
| $L$ | 96 $\mu\text{m}$ | Length of soma |
| $D$ | 96 $\mu\text{m}$ | Diameter of soma |

The conversion between intracellularly injected current values and current density is required by an appropriate scale factor F:

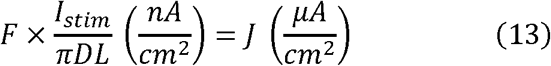

To generate an effective action potential, we first found required current intensity of standard envelope-modulated stimulation that can evoke an action potential in every beat frequency. For example, with regard to carrier frequency of 2 kHz and beat frequency of 100 Hz, the required injected current (*I*_*stim*_ = 200 *nA*) is equivalent to current density (*J* = 690 *µA*/*cm*^2^) calculated from FEM.

### Integration of FEM and neuronal action potential model

In a given divided time, a standard envelope is a superposition of four channels with same amplitude, different frequency (*f*_*c*_ + Δ*f, f*_*c*_) and polarity (positive, negative). Hence, we set the voltage applied to each electrode pair to a value that make the induced current density at the target point close to half of the required current intensity obtained from NEURON. For a given divided time, the total current density at every point is temporally summation of the current densities acting on that divided time.

Following above temporally summation, the induced current densities from FEM were integrated into certain stimulation waveforms. The current waveforms, as an injected current stimulating the soma, was imported into NEURON to calculate membrane potential to determine whether every envelope can evoke an effective action potential. Basically, neural activity at one point is determined by the number of effective action potentials generated in a complete cycle in response to specific stimulation waveform. Finally, the excitable area is defined as the number of action potentials generated in complete cycle is less than half of the maximum value. For neuronal excitability across brain, points were linearly interpolated using MATLAB’s griddata function.

## 3 RESULTS

The optimal stimulation frequency for deep brain stimulation (DBS) is 100-180 Hz (Montgomery, 2017) owing to its potential mechanisms that temporal summation of action potentials can produce the most efficient oscillation. DBS has strict requirements on stimulating frequency.

Therefore, to achieve the best stimulus efficiency, we utilized two carrier frequencies of 2 kHz and 2.1 kHz, resulting in beat frequency of 100 Hz, in accordance with the optimal stimulation frequency for DBS. The frequency and polarity (positive or negative) of scalp-applied current in one cycle (80 ms) are listed in Table S1 and arrangement of multiple electrodes is shown in Fig. S1. The first group of channels (CH 1 - CH 4) and second group of channels (CH 5 – CH 8), with duty cycle of 50 % and complete cycle of 80 ms, formed standard low-frequency envelope in a certain time slot respectively. Therefore, a complete cycle was 80 ms and the frequency of envelope was 100 Hz.

As demonstrated in Fig. 4, the region where neurons generated the most excitability was controlled in a small circle thanks to TDFPMS. With further refinement, the change of number of action potentials along x-axis is shown in Fig. 5. It indicated that deviating 5 mm away from the target location, the number of action potentials dropped instantaneously to less than four (maximum is eight). In summary, the target region was controlled with a circle with radius of 5 mm even though external electrodes were far away 10 cm from the center.

**Figure 4:**
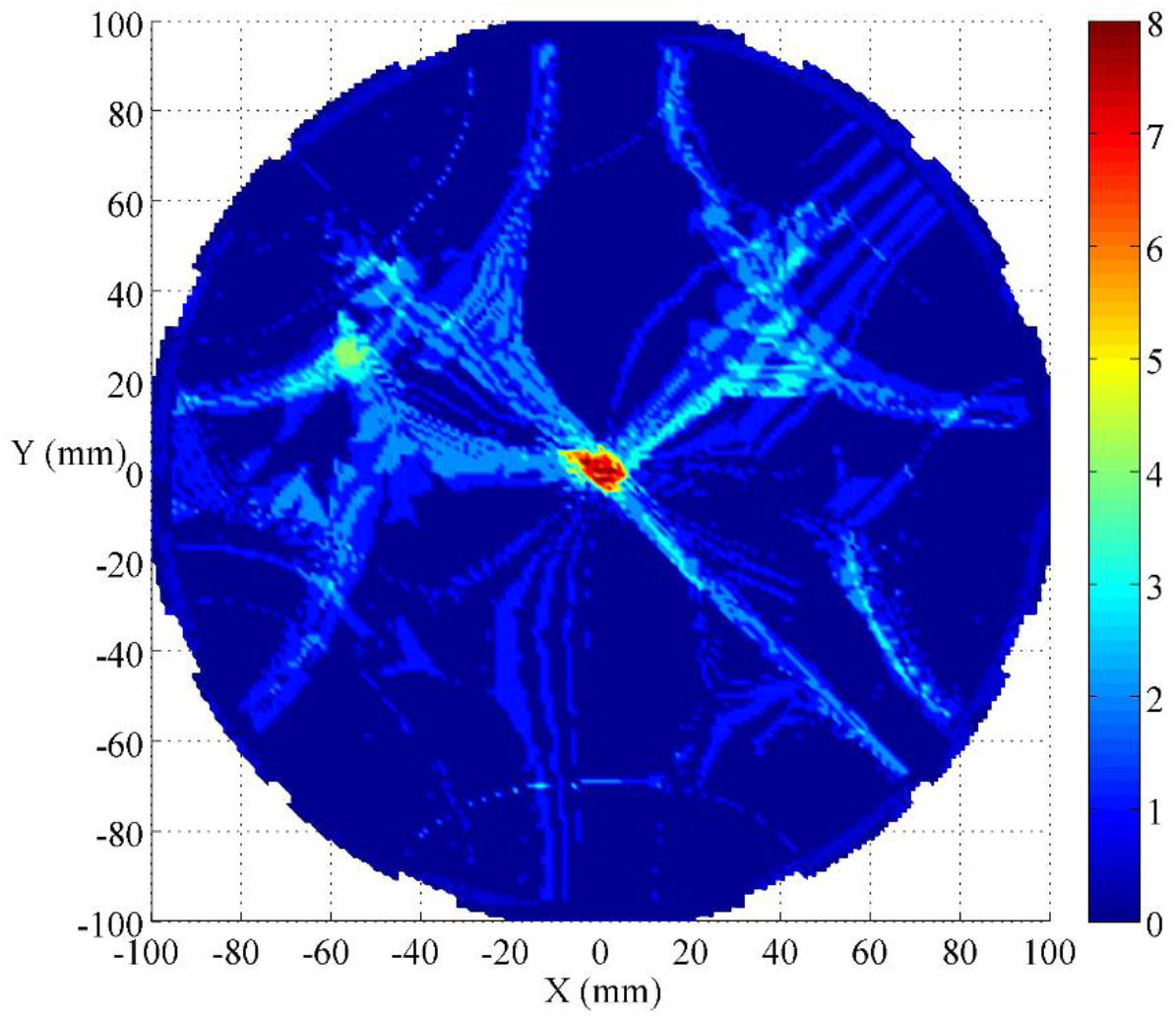
Distribution of neural excitability across the brain when the target location was at center.

**Figure 5:**
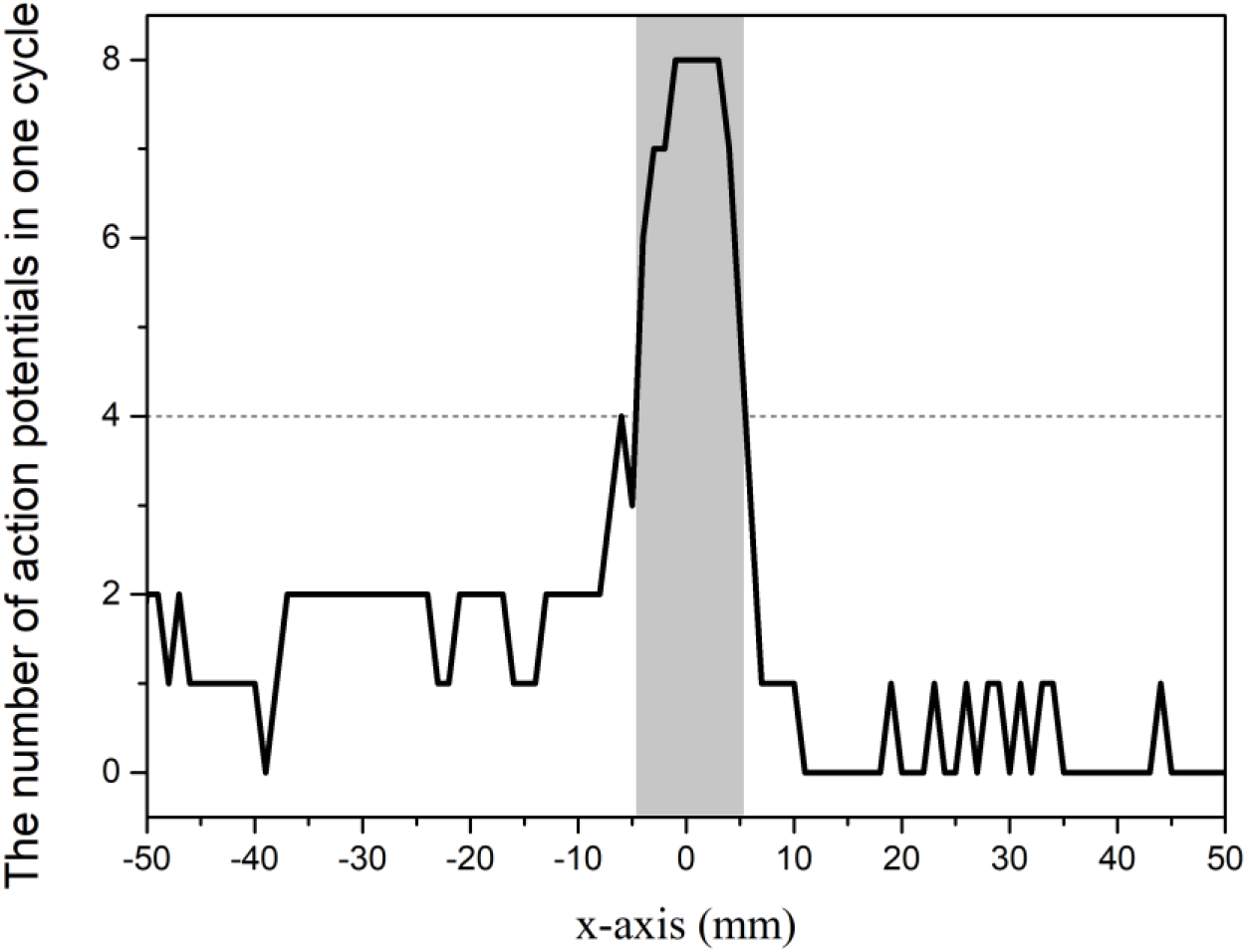
The number of action potentials along x-axis during complete cycle (80 ms). The maximum number of action potentials in target region was eight while deviating 5 mm from the target region dropped to less than four.

To demonstrate our method’s steerability, we set the target region in off-center area with coordinate (15 mm, −5 mm), consistent with the imaging position of the subthalamic nuclears (STN) on the axial plane (Hamid et al., 2005). It just adjusted each channel’s outputting amplitude to a certain value that current densities were equal in the STN, without electrode movement. As demonstrated in Fig. 6, the most excitable region was STN.

**Figure 6:**
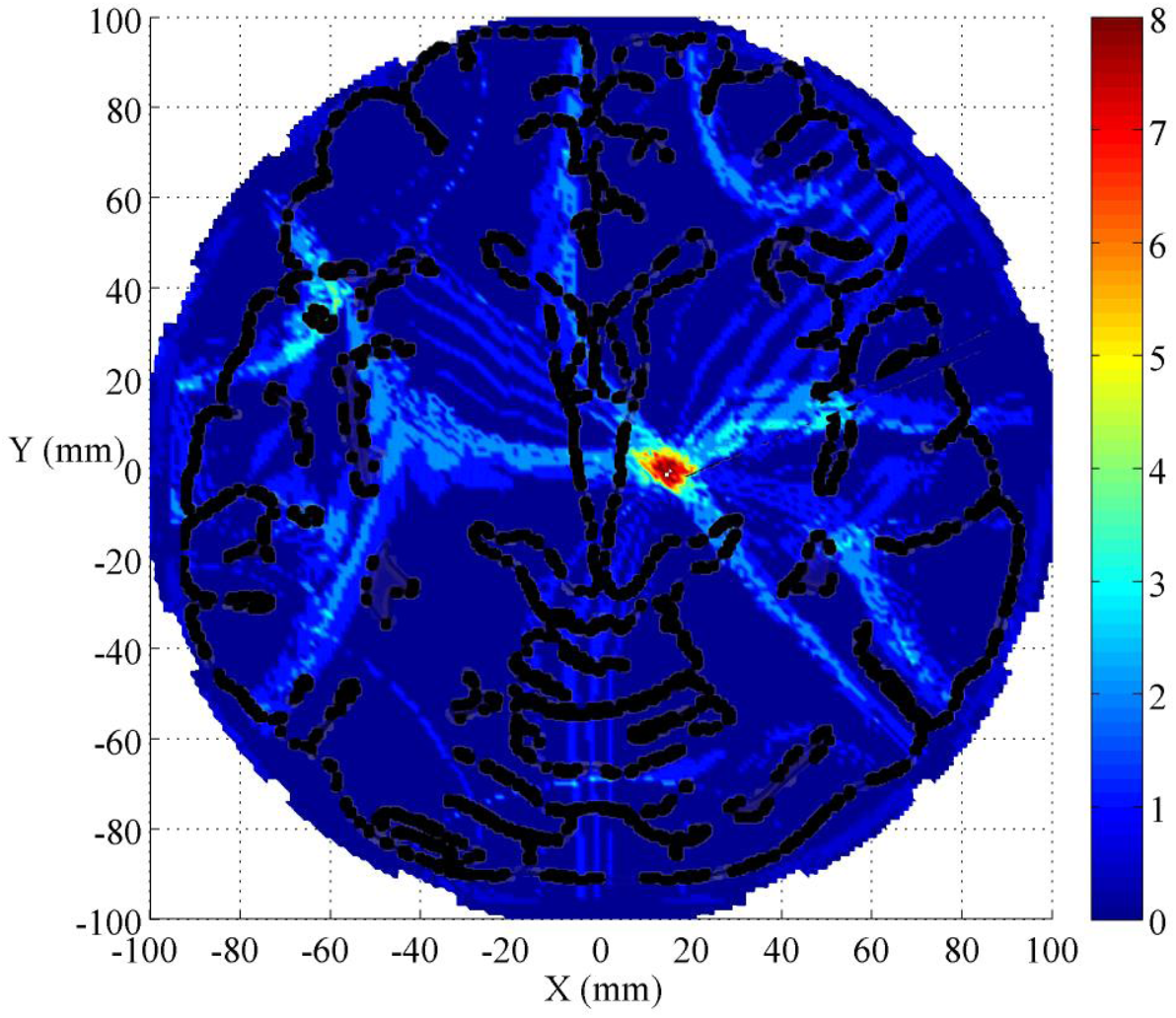
Distribution of neural excitability across the brain when target location is off center: STN.

Within the target stimulation region, the induced current densities by each electrode pair were approximately equal. As shown in Fig. 7, during every 10 ms (100 Hz), the soma evoked an effective action potential in response to standard low frequency envelope, modulated at the frequency of 100 Hz. In a complete cycle, the effect of stimulation waveform was approximately equivalent to continuous 100 Hz sinusoidal wave and elicit optimal stimulus efficiency.

**Figure 7:**
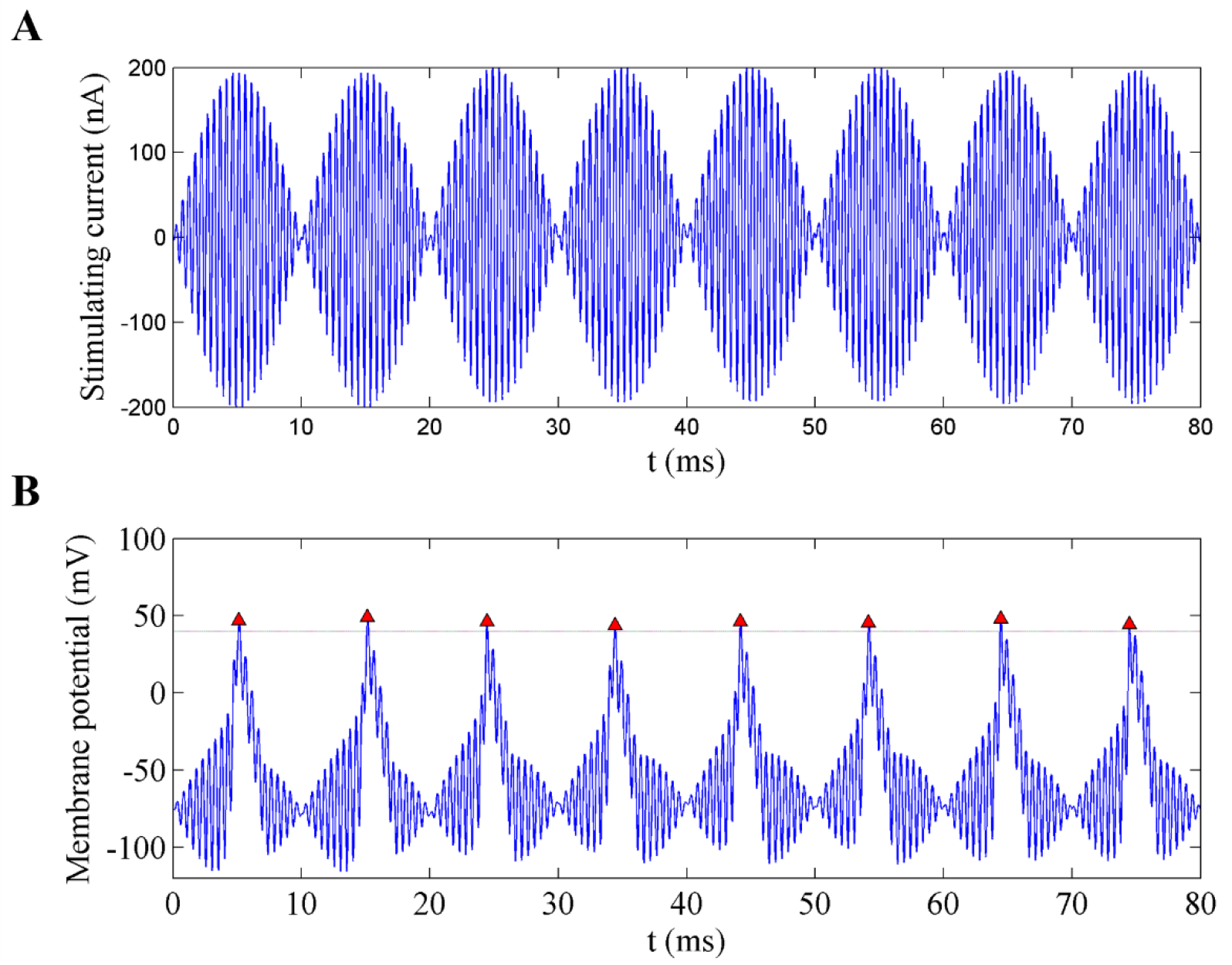
Stimulation waveform and membrane potentials in target region. (A) Temporally summation of stimulation waveform, with envelope frequency of 100 Hz and carrier frequency of 2 kHz. (B) The membrane potential of soma in NEURON corresponding to above stimulation. The red marker indicates an effective action potential.

Deviating from target location, as illustrated in Fig. 8, the current densities generated by various channels were no longer equal, resulting in not effective stimulation. Inevitably, in the switch of time division, it may induce an action potential. However, it can be neglected because temporal summation of action potential cannot be established. Firing rate (or the number of action potentials in complete cycle) and stimulus efficiency were quite low compared with the central area.

**Figure 8:**
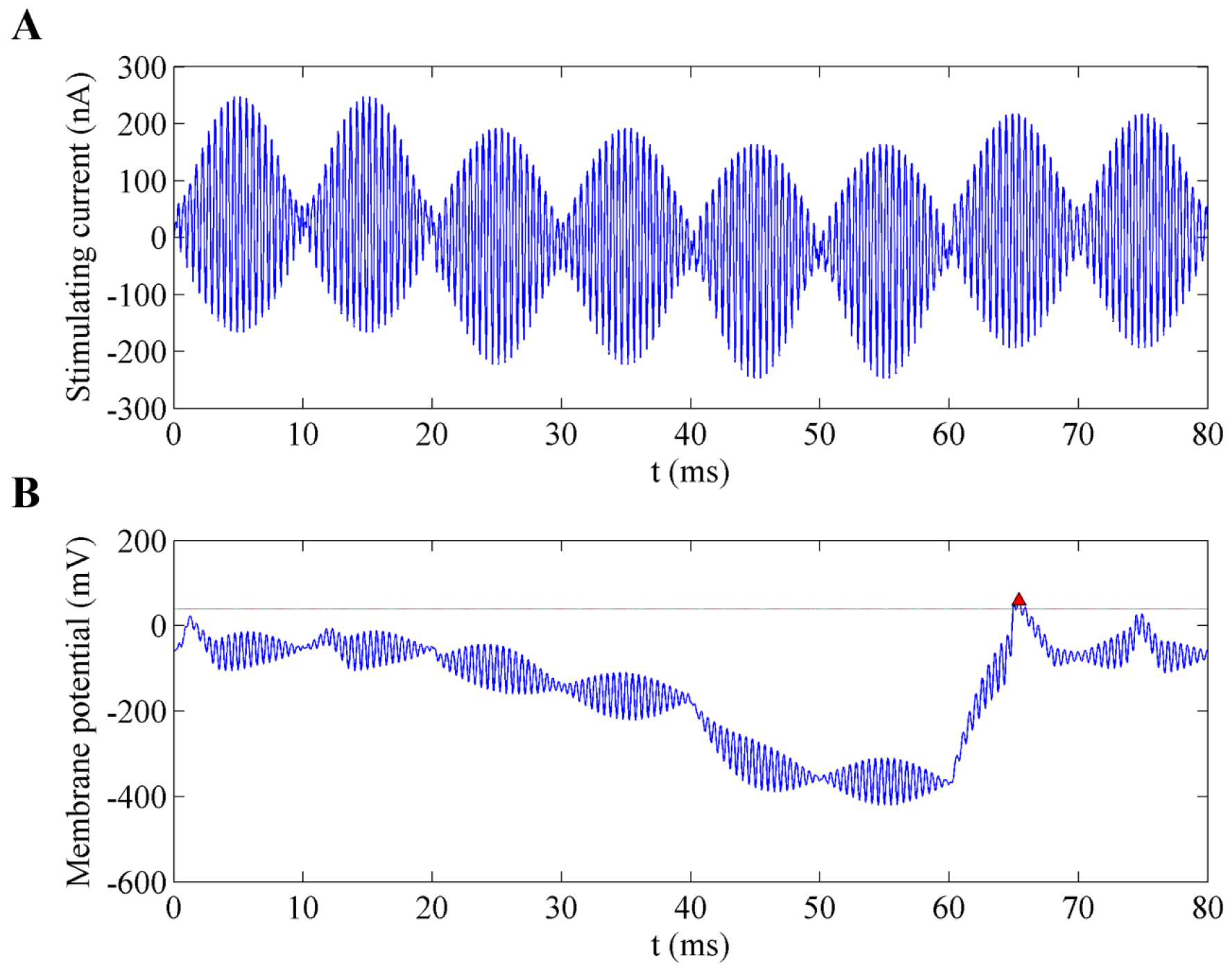
Stimulating waveform and membrane potentials in the region deviated from target region. (A) Temporally summation of stimulating waveform, with different induced current densities: *I*_*stim* 1_ = 118.80 *nA I*_*stim* 2_ = 91.88 *nA I*_*stim* 3_ = 74.70 *nA I*_*stim* 4_ = 131.44 *nA I*_*stim* 5_ = 90.06 *nA I*_*stim* 6_ = 119.91 *nA I*_*stim* 7_ = 120.70 *nA I*_*stim* 8_ = 75.31 *nA*. (B) The membrane potential of soma in NEURON corresponding to above stimulation waveform. The red marker indicates an effective action potential and the membrane potential is above 30 mV.

## 4 DISCUSSION

We have described here a noninvasive TDFPMS method that can effectively stimulate deep neurons with high spatial precision (mm-level) via finite-element and action potential modelling. Additionally, we verified the correctness of this method through measuring distribution of stimulation waveforms in saline solution. High focality is achieved by multiple electrodes applying positive and negative polarity outputting separately, time-division and different kilohertz-frequency. The excitable region is not only determined by current density just like traditional method, but specific characteristic of waveform should be taken into consideration.

It is important to note that TDFPMS improving spatial focality is an adaptation of the concept of “time division multiple access” in communication field. The complete stimulating cycle is divided into several time slots. One group of electrodes, acting on that divided time, control one target location where neurons are most excitable. The other group of electrodes control the other target region during the other divided time. Therefore, in the complete cycle, standard envelope stimulation only exists in the intersection of above target region where elicit the most action potentials and excitability. Deviating from the target location, not all envelope can evoke firing event, resulting in reduction of neural excitability. At the same time, electrical stimulation has its own optimal frequency, below which the stimulus efficiency is greatly reduced or without therapeutic effect. Therefore, although there are a few action potentials in the edge region, it can be neglected. Choosing the best envelope frequency, the same as the optimal stimulation frequency, our time-division method maximizes the activity of neurons in the target location.

In future studies, to improve spatial resolution and practical operability of electrodes, we will get MRI scans to create a more realistic human model with accurate mesh type optimize arrangement of electrodes and number of time division. Furthermore, we will introduce living neural experiments to further verify practicability of our method and its clinical therapeutic effect.

## 5 CONCLUSION

In conclusion, we propose noninvasive method can stimulate neurons at depth with high spatial resolution. We demonstrate its mm-level spatial resolution by finite-element analysis and action potential modeling. Our proposed method that utilize time-division, frequency & polarity modulation synthesis (TDFPMS) will provide a basic scheme for noninvasive deep neurostimulation.

